# CycDelta: Multimodal Residue-Graph Learning of Permeability Changes in Cyclic Peptides

**DOI:** 10.64898/2026.09.18.752569

**Authors:** Jie Qian, Yu Zhou, Yaning Cui, Zhifeng Gao, Lingyun Wu, Sean Li, Maohua Yang, Dongdong Wang

## Abstract

Membrane permeability remains a major challenge in cyclic-peptide drug discovery, where optimization often depends on estimating changes relative to measured reference compounds. Existing predictors mainly model absolute permeability, while heterogeneity across literature sources complicates pooled training and comparative evaluation. We present CycDelta, a multimodal residue-graph model trained directly on within-source permeability differences. CycDelta combines directed message passing with pretrained Uni-Mol monomer embeddings, physicochemical descriptors and assay information in a shared encoder for reference and query peptides. To evaluate this comparative formulation under controlled conditions, we establish CycDeltaBench, comprising in-distribution, scaffold-based out-of-distribution, external and zero-shot unseen-assay tests. On the out-of-distribution task, CycDelta achieved Pearson 0.69 versus 0.46 for the strongest baseline. It also led all evaluated methods on two external data sets and achieved zero-shot Spearman 0.70 versus 0.32 for the strongest baseline on an unseen assay. Residue-level attribution relates predictions to monomer contributions, while gradient-guided screening supports prioritization of permeability-enhancing analogue hypotheses. These results position CycDelta as a transferable and interpretable framework for comparative permeability prediction and cyclic-peptide optimization.

## 1 Introduction

Cyclic peptides occupy a useful region between small molecules and biologics [1–3]. Their extended interaction surfaces can engage difficult targets, while conformational restriction can provide affinity, selectivity and, in favourable cases, oral exposure or intracellular access [3, 4]. Membrane permeability remains a central obstacle and has motivated experimental design rules, permeability databases and machine-learning models for cyclic peptides [5–8].

Experimentally, cyclic-peptide permeability is characterized using several complementary assay formats. PAMPA estimates passive diffusion across an artificial lipid membrane in a cell-free system [9]. Caco-2 monolayers model the intestinal epithelial barrier and can reflect both passive transport and transporter effects [10]; MDCK monolayers provide a comparatively rapid cellular permeability screen [11]; and low-efflux RRCK cells are designed to reduce transporter-mediated efflux and more closely emphasize passive permeability [12]. Because these assays capture related but non-identical transport processes, their numerical outputs are not directly interchangeable [7, 13]. Published cyclic-peptide predictors therefore commonly define assay-specific training data rather than pooling measurements from different assay formats as equivalent labels [8, 14–17]. However, fixing the assay does not remove variability across sources (the publications or reports providing the measurements). In CycPeptMPDB, where permeability is reported as Log*P*_exp_, the base-10 logarithm of the permeability coefficient in cm s^−1^ [7], we identified 21 groups comprising 57 PAMPA measurements of the same structure across different sources. Their mean within-group sample standard deviation was 0.63 Log*P*_exp_ units, corresponding to an approximately 10^0.63^ = 4.3-fold factor in permeability. For comparison, 414 groups comprising 832 measurements of the same structure and source across different assay methods had a mean standard deviation of 0.55 units, or approximately 3.5-fold. Cross-source variability within PAMPA was therefore comparable in magnitude to variability across assay formats and can introduce substantial label noise when measurements from heterogeneous literature sources are pooled. The cross-source comparisons are shown in Supplementary Fig. S1.

Most computational formulations predict an absolute assay value for one peptide at a time. Although scientifically useful, this endpoint does not fully match a common medicinal-chemistry decision: given a measured lead or reference peptide, which candidate is expected to improve permeability, and by how much? Medicinal-chemistry programmes often prioritize analogues by asking whether a structural modification improves a property relative to a measured lead or reference peptide, rather than by predicting the absolute property of each analogue independently. This comparative perspective is related to matched molecular pair analysis [18, 19]. For a query *q* and reference *r*, the corresponding endpoint is Δ*P*_*q,r*_ = *P*_*q*_−*P*_*r*_. When both measurements come from the same source, subtraction can partially control source-specific effects and focus evaluation on the permeability change associated with the molecular difference.

The release of CycPeptMPDB enabled predictors that progressively incorporate sequence, graph, physico-chemical, image and multimodal representations. Representative examples include Multi_CycGT, which combines SMILES, atom-level graph and descriptor features [14]; CycPeptMP, which integrates atom-, monomer- and peptide-level information [15]; MultiCycPermea, which aligns SMILES and molecular-image encoders [8]; and SinCAA, which learns representations tailored to non-canonical amino acids [20]. Related architectures use pretrained sequence encoders and multimodal feature fusion [16, 17]. A systematic comparison of 13 fingerprint-, string-, graph- and image-based methods identified DMPNN as a strong general baseline and reported substantial losses under scaffold splitting [21]. More recent studies have further explored applicability-domain calibration, generative optimization, monomer-aware and multimodal representation learning, and conformationally informed prediction [22–27].

Despite this progress, comparison among published models remains complicated by inconsistent curation, assay selection and train–test splitting [13, 28]. Random record-level splits can distribute related scaffolds across training and test sets, leading to optimistic estimates of structural generalization [29–32], while models evaluated on different collections or endpoints cannot be ranked reliably. However, to our knowledge, no standardized benchmark has systematically evaluated prediction of permeability differences between related cyclic peptides measured within the same source. Addressing this gap requires fixed records, partitions, pair construction, reference selection and metrics, together with both in-distribution (ID) and scaffold-level out-of-distribution (OD) evaluation.

Here we present CycDelta, a multimodal residue-graph model designed to learn permeability changes directly between a query peptide and a measured reference from the same source. CycDelta integrates directed message passing, pretrained Uni-Mol monomer embeddings, RDKit descriptors and assay information in a shared molecular encoder. To evaluate this comparative formulation consistently, we establish CycDeltaBench, which fixes the records, query–reference construction and evaluation protocol across ID, scaffold-OD, external and unseen-assay settings. The model is not restricted to predefined modification types and is designed to support both permeability prediction and analogue prioritization.

This work makes three main contributions: (i) it introduces CycDelta, a multimodal residue-graph model trained directly on within-source permeability differences and achieving leading performance on the primary OD task and across the external and unseen-assay evaluations; (ii) it establishes CycDeltaBench, which defines the pairwise task and organizes the available data into fixed ID and scaffold-clustered OD partitions, external tests on the Merz and Faris data, and a zero-shot unseen-assay test on the Nielsen data; and (iii) it incorporates residue-level attribution and gradient-guided monomer substitution to support interpretation and cyclic-peptide permeability optimization. Together, these components connect a task-aligned predictive model with controlled evaluation and practical analogue prioritization.

## 2 Task formulation and evaluation protocol

### 2.1 Within-source pairwise endpoint

For peptide *i* measured in source *s*, we write

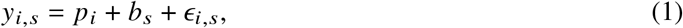

where *p*_*i*_ is the molecular contribution of interest, *b*_*s*_ is an additive source-associated term and *ϵ*_*i,s*_ is residual error. For query *q* and reference *r* from the same source, the benchmark target is

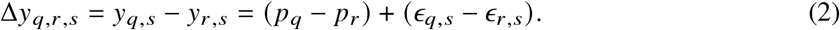

The additive source term cancels under this model. This argument motivates within-source target construction but does not imply that all inter-source effects are additive or that two records from one source are experimentally identical in every respect.

Each source must contain at least two eligible records in a split. One reference peptide is sampled independently per source; the other eligible records act as queries. References are resampled during training to avoid dependence on one anchor. Every directed pair (*q, r*, Δ*y*) is augmented with (*r, q*, −Δ*y*). Validation and reported sensitivity use ten fixed reference-selection seeds.

### 2.2 Data curation

Development records were curated from CycPeptMPDB v1.2 [7], following the general principle that molecular structures and assay annotations should be checked before model development [13, 28]. Records with permeability equal to −10 were removed. We also removed data from Koch et al. study [33], because the SMILES annotations for these records in CycPeptMPDB were found to contain problems. The resulting development collection contained 8,303 records.

The ID split used random record-level allocation. The OD split exactly reproduced the scaffold-based partitioning procedure used by MultiCycPermea [8], grouping peptides according to scaffold similarity and assigning complete scaffold groups to the training, validation and test sets. Scaffold-based partitions are commonly used to provide a more demanding test of chemical generalization than random record splits [29, 30]. Table 1 summarizes the development partitions and external evaluation sets.

**Table 1.**
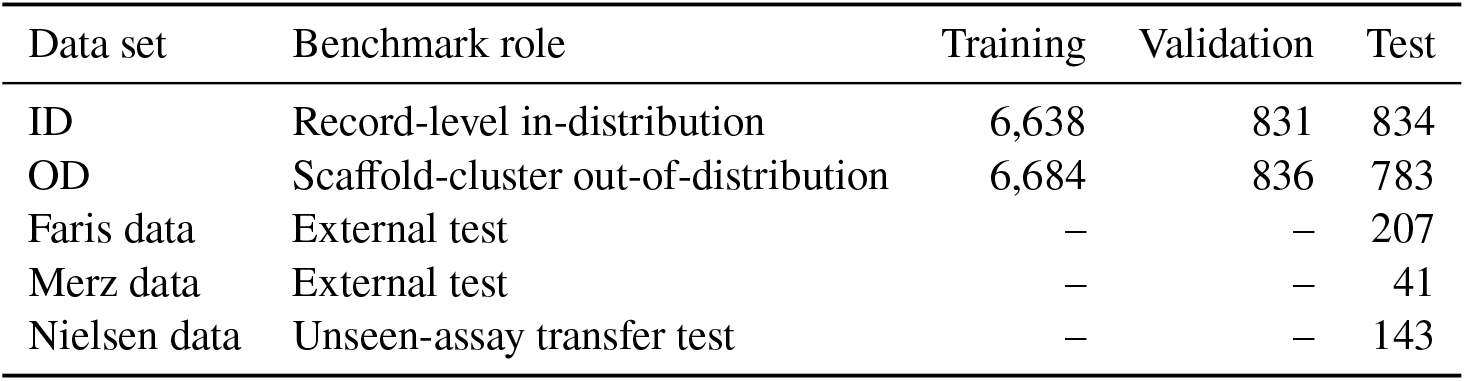
CycDeltaBench development and external evaluation sets. Counts refer to records. External sets were used only for testing.

| Data set | Benchmark role | Training | Validation | Test |
| --- | --- | --- | --- | --- |
| ID | Record-level in-distribution | 6,638 | 831 | 834 |
| OD | Scaffold-cluster out-of-distribution | 6,684 | 836 | 783 |
| Faris data | External test | — | — | 207 |
| Merz data | External test | — | — | 41 |
| Nielsen data | Unseen-assay transfer test | — | — | 143 |

Two literature-derived data sets were reserved for external evaluation. The Faris data comprise 207 records reported by Faris et al. [34], and the Merz data comprise 41 records reported by Merz et al. [35]; both report permeability as Log*P*_exp_. The Nielsen data contain 143 cyclic peptides evaluated by the chloroalkane penetration assay (CAPA), with cellular penetration reported as HaloTag occupancy (% HaloTag reaction) after a 4-h incubation with 1 *μ*M compound [36, 37]. CAPA was absent from the four assay categories used during model training. Because the Nielsen endpoint differs from Log*P*_exp_ in both physical meaning and numerical scale, transfer performance on this data set is reported using Spearman correlation, which evaluates rank agreement without assuming scale equivalence.

The Merz, Faris and Nielsen data shared no scaffold cluster with the internal development collection under the clustering procedure used here. In other words, none of the three evaluation sets was assigned to a scaffold cluster represented during model development. Figure 1 provides a qualitative two-dimensional view of their chemical-space distributions. Cluster assignments, rather than apparent separation in the projection, defined scaffold overlap.

**Figure 1.**
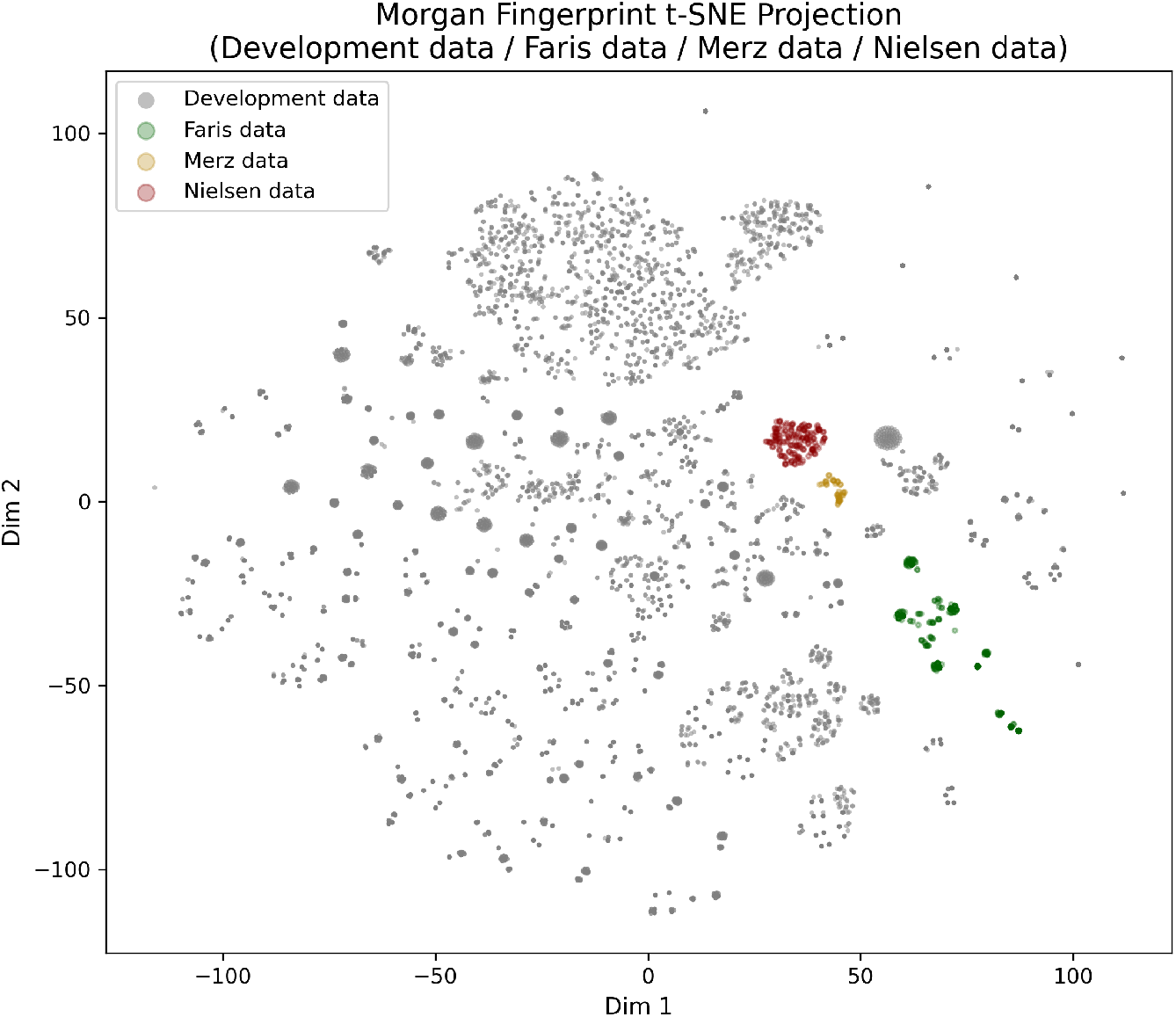
Chemical-space distributions of the development and three external evaluation sets. A t-SNE projection of Morgan fingerprints [38] is shown for the internal development data and the Merz, Faris and Nielsen data sets.

### 2.3 Evaluation suites and metrics

CycDeltaBench contains five complementary suites:

1. ID within-source Δ prediction;
2. scaffold-clustered OD within-source Δ prediction, designated the primary internal leaderboard;
3. ID and OD absolute-value prediction as contextual comparisons;
4. external pairwise evaluation on the Merz and Faris data; and
5. zero-shot assay transfer on the Nielsen data.

Within-source evaluation measures agreement with Δ*y*. For a pairwise model, absolute predictions are reconstructed from one shared measured anchor:

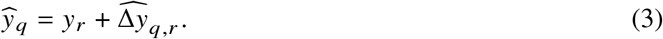

Pearson correlation measures linear agreement, Spearman correlation measures rank agreement, and mean absolute error (MAE) measures absolute deviation. Higher correlations and lower MAE are better. Values after “±” denote standard deviation over ten reference selections and describe anchor sensitivity.

### 2.4 Compared methods

The comparison includes Multi_CycGT [14], graph convolutional networks (GCN) [39], graph attention networks (GAT) [40], Morgan-fingerprint regression [38], a Gaussian process using Tanimoto similarity [25], SinCAA with its pretrained non-canonical amino-acid representations and downstream GIN predictor [20], MultiCycPermea [8], DMPNN, which was identified as the strongest overall method across the benchmark tasks evaluated by Liu et al. [21], and CycDelta. All methods use the same records, partitions and evaluation targets. The benchmark distinguishes target definition from architecture: conventional absolute predictors are converted to pairwise changes by subtracting their predictions for the query and reference, whereas CycDelta is trained directly on within-source differences.

Head-to-head evaluation was restricted to methods whose inference pipelines could be executed or faithfully reproduced under the fixed CycDeltaBench records, partitions and targets. Other published systems were retained in the related-work discussion but not ranked when an end-to-end implementation or trained model was unavailable, when required inputs could not be generated reproducibly under the benchmark protocol, or when the reported endpoint was not compatible with the present prediction tasks.

## 3 CycDelta model

### 3.1 Cyclic-peptide residue graphs

SMILES strings were segmented at macrocyclic amide bonds to obtain residue fragments. Canonical rotation removed the arbitrary start position of a cyclic sequence. Residues matching the 20 natural amino acids were assigned their identities; all other monomers were assigned an unknown-residue class. A peptide with *L* residues was represented as a complete directed graph *G* = (*V, E*), where *V* = {1, …, *L*} and

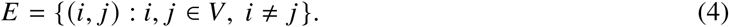

Each directed edge has a scalar structural attribute *a*_*ij*_ : *a*_*ij*_ = 1 for adjacent backbone residues and the ring-closing pair, and *a*_*ij*_ = 0 otherwise. Thus, non-local residue pairs can exchange messages while backbone connectivity remains explicitly marked.

### 3.2 Multimodal residue features

The basic feature vector for residue *i* is

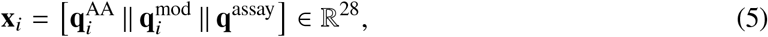

where 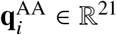 is a natural/unknown residue one-hot vector, 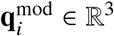 encodes natural-residue status, D stereochemistry and methylation, and **q**^assay^ ∈ R^4^ encodes PAMPA, CACO2, MDCK or RRCK and is broadcast to all residues.

Each residue additionally receives a 512-dimensional Uni-Mol embedding [41]. To express the pretrained representation relative to a common monomer, we use

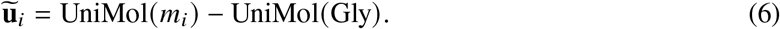

A 31-dimensional RDKit descriptor vector **d**_*i*_ [42], containing size, polarity, lipophilicity and shape descriptors, is standardized using training-set means and standard deviations; non-finite values are replaced by the corresponding training mean before standardization.

This multimodal representation is particularly well suited to non-canonical amino acids: although they share the unknown-residue identity class, their monomer-specific Uni-Mol embeddings, modification flags and RDKit descriptors preserve differences in three-dimensional and physicochemical features rather than collapsing all non-canonical residues into a single undifferentiated token.

The three modalities are projected separately and fused:

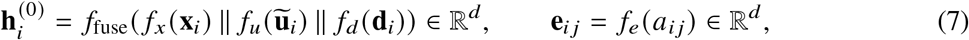

where *d* = 128 and all *f* functions are multilayer perceptrons with LeakyReLU activations.

### 3.3 D-MPNN encoding and delta regression

Figure 2 summarizes the architecture. The fused residue and edge features are processed by a standard directed message-passing neural network (D-MPNN) module [43]. The resulting residue states are sum-pooled to obtain a fixed-dimensional graph representation **g**(*G*).

**Figure 2.**
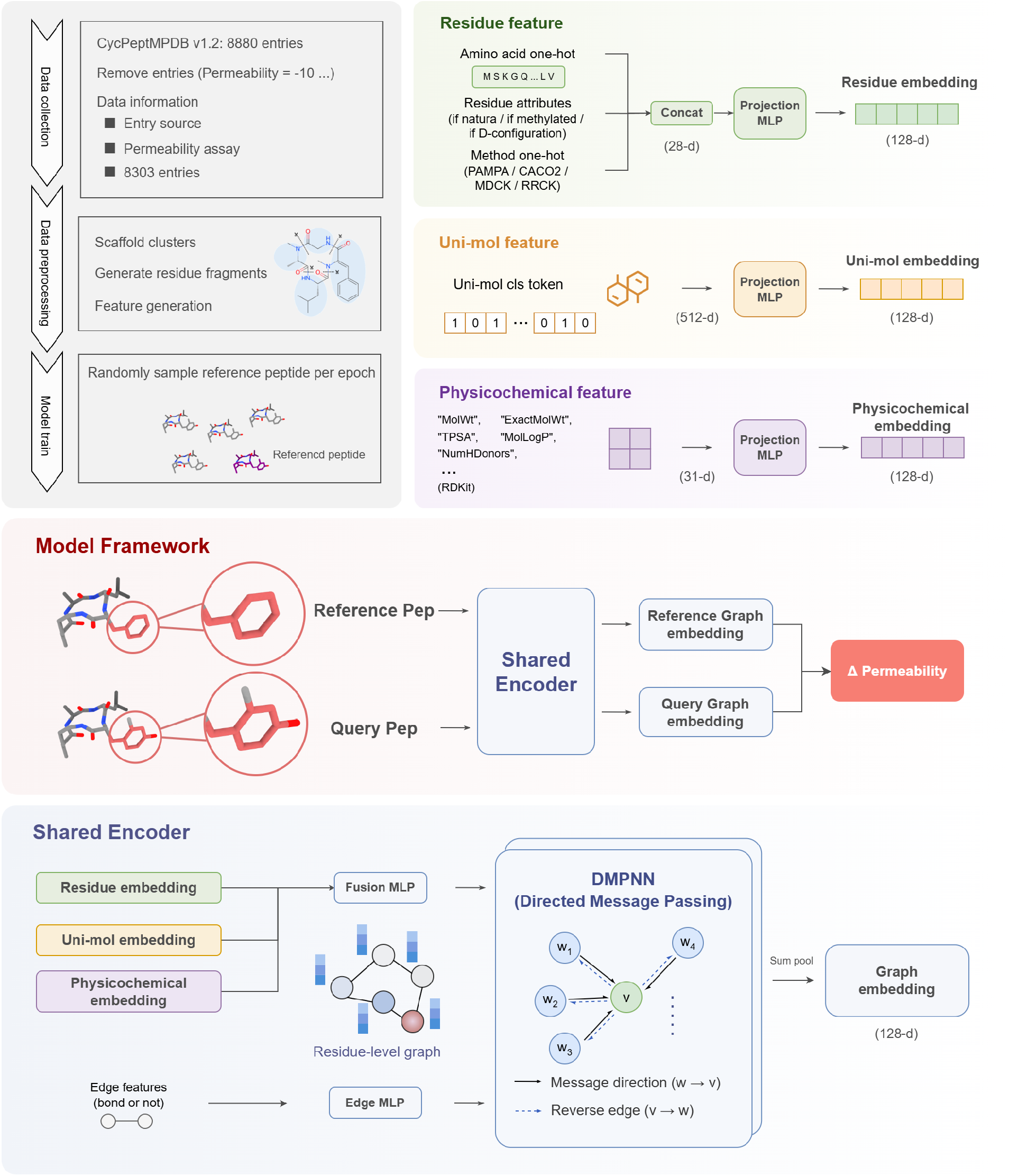
Overview of CycDelta. Cyclic peptides are segmented into residues and represented using residue identity and modification flags, assay identity, glycine-relative Uni-Mol embeddings and standardized RDKit descriptors. A shared D-MPNN encodes the reference and query peptides. Their graph embeddings are subtracted and regressed to the permeability difference. The lower panel details directed-edge message passing and graph readout.

The reference and query graphs are encoded by the same network. Their embedding difference is passed to the output multilayer perceptron:

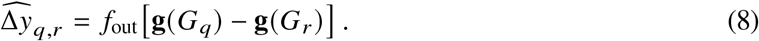

The resulting architecture is a shared-weight delta regressor trained specifically for within-source permeability differences.

### 3.4 Training

At the beginning of each training epoch, one reference peptide was randomly resampled for every eligible source, and the remaining peptides from that source were used as queries. During model fitting, references were selected exclusively from the training partition; no validation or test records were used to construct training pairs, thereby preventing data leakage. Every resulting pair (*q, r*, Δ*y*) was augmented by its reverse (*r, q*, −Δ*y*). For *N* directed training pairs, model parameters *θ* were optimized with mean-squared error:

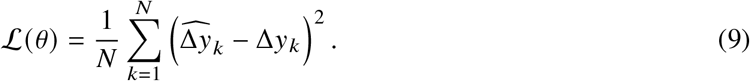

The default configuration used two D-MPNN message-passing layers and Adam with learning rate 10^−4^, batch size 32 after bidirectional augmentation, dropout 0.25, at most 2,000 epochs.

For few-shot fine-tuning on the Nielsen data, all feature projections and the D-MPNN encoder were frozen, graph embeddings were precomputed, and only *f*_out_ was optimized with the delta MSE objective and bidirectional augmentation. Fractions from 5% to 80% of the 143 CAPA-labelled peptides were sampled in ten seeded repeats; the selected fraction was used to fine-tune the output head and the remaining peptides formed the test set. No separate validation set was used to choose the epoch. Epoch-wise comparisons are consequently descriptive and test-informed, and the few-shot experiment is kept distinct from the prespecified zero-shot benchmark.

## 4 Model performance

### 4.1 OD setting performance

To assess whether the models generalize beyond scaffold groups represented during training, we first evaluated them under the scaffold-OD setting. Table 2 presents the within-source leaderboard together with the absolute-value context. CycDelta achieved Pearson *r* = 0.69, Spearman ρ = 0.67 and MAE = 0.74 on within-source differences. Morgan FP provided the second-highest correlations, 0.46 and 0.44, giving CycDelta margins of 0.23 for both metrics. Multi_CycGT obtained the next-lowest MAE of 0.75. Thus, CycDelta’s clearest OD advantage lies in preserving relative ordering and covariation after source-associated offsets are removed. Representative observed-versus-predicted scatter and regression plots for CycDelta at reference-selection seed 0 are shown in Supplementary Fig. S2.

**Table 2.** OD benchmark leaderboard. Higher Pearson and Spearman and lower MAE are better. “±” denotes standard deviation over ten reference selections. Bold and underline mark the best and second-best values, respectively.

| Target | Method | Pearson | Spearman | MAE |
| --- | --- | --- | --- | --- |
| Within-source $\Delta$ | Multi_CycGT | 0.34±0.08 | 0.33±0.08 | <u>0.75±0.09</u> |
|  | GCN | 0.09±0.25 | 0.07±0.28 | 1.08±0.26 |
|  | GAT | 0.18±0.20 | 0.14±0.21 | 1.07±0.26 |
|  | Morgan FP | <u>0.46±0.22</u> | <u>0.44±0.22</u> | 0.93±0.19 |
|  | GP + Tanimoto | 0.40±0.12 | 0.38±0.12 | 0.84±0.09 |
|  | SinCAA | 0.37±0.06 | 0.38±0.08 | 0.78±0.07 |
|  | MultiCycPermea | 0.34±0.15 | 0.32±0.16 | 0.77±0.12 |
|  | DMPNN | 0.36±0.05 | 0.36±0.08 | 0.79±0.09 |
|  | CycDelta | **0.69±0.11** | **0.67±0.11** | **0.74±0.14** |
| Absolute value | Multi_CycGT | 0.23 | 0.22 | 0.79 |
|  | GCN | 0.23 | 0.31 | 0.80 |
|  | GAT | 0.31 | 0.33 | 0.86 |
|  | Morgan FP | 0.44 | 0.51 | 0.78 |
|  | GP + Tanimoto | 0.46 | 0.43 | 0.81 |
|  | SinCAA | 0.46 | <u>0.54</u> | <u>0.76</u> |
|  | MultiCycPermea | 0.48 | 0.49 | 1.01 |
|  | DMPNN | <u>0.51</u> | **0.55** | **0.72** |
|  | CycDelta | **0.53±0.05** | 0.48±0.05 | 0.88±0.20 |

The absolute-value view was mixed. CycDelta had the highest Pearson correlation at 0.53, only 0.02 above DMPNN. DMPNN had the highest Spearman correlation (0.55) and lowest MAE (0.72), followed by SinCAA at 0.54 and 0.76, whereas CycDelta obtained 0.48 and 0.88. The benchmark therefore does not support a claim that CycDelta is uniformly superior for absolute permeability prediction.

Notably, several of the strongest conventional baselines on OD absolute-value prediction lost correlation after evaluation was changed to within-source differences. DMPNN decreased from Pearson/Spearman 0.51/0.55 to 0.36/0.36, MultiCycPermea from 0.48/0.49 to 0.34/0.32, GP + Tanimoto from 0.46/0.43 to 0.40/0.38, and SinCAA from 0.46/0.54 to 0.37/0.38. One possible explanation is that their absolute predictions partly exploit systematic differences among literature sources. Within-source subtraction removes source-level offsets and weakens that signal, forcing the model to resolve the finer relationship between structural changes in a peptide pair and the associated permeability change. A related hypothesis is that literature sources occupy different scaffold distributions, allowing an absolute model to learn the broad property range associated with a scaffold family rather than the detailed mapping from peptide modification to property difference.

### 4.2 ID setting performance

To establish performance under standard record-level in-distribution splitting, we next evaluated all methods in the ID setting. Complete results are provided in Table 3. On the ID within-source task, CycDelta reached Pearson 0.89 and Spearman 0.87, exceeding the next-highest values by 0.16 and 0.21. Its MAE of 0.56 did not lead the benchmark; GCN achieved 0.48. In the ID absolute-value setting, CycDelta obtained 0.64/0.63 correlations and MAE 0.59, while DMPNN and GCN led individual metrics. The corresponding seed-0 observed-versus-predicted scatter and regression plots are provided in Supplementary Fig. S3.

**Table 3.** ID benchmarks under absolute-value and within-source Δ evaluation. Higher correlations and lower MAE are better.

| Target | Method | Pearson | Spearman | MAE |
| --- | --- | --- | --- | --- |
| Within-source $\Delta$ | Multi_CycGT | 0.60±0.13 | 0.60±0.14 | 0.52±0.08 |
|  | GCN | <u>0.73±0.12</u> | <u>0.66±0.13</u> | **0.48±0.09** |
| | GAT | 0.71±0.13 | 0.63±0.15 | 0.55±0.08 |
| | Morgan FP | 0.70±0.11 | 0.65±0.12 | 0.55±0.08 |
| | GP + Tanimoto | 0.60±0.08 | 0.64±0.07 | 0.54±0.06 |
| | SinCAA | 0.61±0.11 | 0.59±0.13 | 0.51±0.08 |
| | MultiCycPermea | 0.57±0.12 | 0.55±0.13 | 0.53±0.06 |
| | DMPNN | 0.62±0.08 | 0.61±0.09 | <u>0.49±0.08</u> |
| | CycDelta | **0.89±0.03** | **0.87±0.04** | 0.56±0.08 |
| Absolute value | Multi_CycGT | 0.60 | 0.60 | 0.48 |
|  | GCN | <u>0.67</u> | **0.66** | **0.41** |
|  | GAT | 0.66 | 0.64 | <u>0.43</u> |
|  | Morgan FP | 0.60 | 0.57 | 0.45 |
|  | GP + Tanimoto | 0.51 | 0.55 | 0.52 |
|  | SinCAA | 0.66 | 0.62 | **0.41** |
|  | MultiCycPermea | 0.63 | 0.59 | 0.53 |
|  | DMPNN | **0.69** | <u>0.65</u> | **0.41** |
| | CycDelta | 0.64±0.02 | 0.63±0.02 | 0.59±0.21 |

CycDelta retained competitive absolute-value correlations in the ID setting while establishing a substantially larger lead on within-source Δ prediction. Unlike the OD results, most baseline models did not show a pronounced decrease when their absolute predictions were converted to within-source differences. DMPNN and MultiCycPermea showed only modest declines, while the decrease for GAT was limited to 0.01 in Spearman correlation. One possible explanation is that random ID splitting preserves related scaffolds and source-associated property distributions across the training and test sets, allowing absolute predictors to retain useful relative information after subtraction. Under the scaffold-OD split, this overlap is reduced, and resolving permeability differences between unseen scaffold groups becomes more dependent on learning a transferable mapping between structural variation and property change. CycDelta’s strong ID Δ performance, together with its competitive absolute-value correlations, therefore supports direct within-source training without sacrificing its ability to model the broader permeability trend.

The ID results also separate ranking quality from numerical calibration. CycDelta produced the strongest ordering and linear association for within-source changes, but its MAE was 0.08 higher than the best value. This pattern suggests that the model captures which peptide is expected to be more permeable more reliably than the exact magnitude of the change. The smaller ID–OD drop for CycDelta correlations relative to several baselines is encouraging for scaffold transfer, although the larger OD reference-selection standard deviations indicate that the chosen anchor remains a source of uncertainty.

### 4.3 External test

To test transfer beyond the internal benchmark, we applied models trained under the scaffold-OD setting to two independent literature data sets. Figure 3a,b compares Spearman correlation on the two external tests; Pearson values are reported in the text for completeness. On the Merz data (*N* = 41), CycDelta achieved Pearson correlation 0.74 and Spearman correlation 0.77. The strongest competing values were 0.57 for both metrics, obtained by Multi_CycGT and SinCAA, respectively. On the larger Faris data set (*N* = 207), CycDelta achieved 0.42 for both correlations, compared with 0.30/0.33 for Multi_CycGT; SinCAA reached 0.22/0.28. The two external evaluations support transfer across literature sources, although the smaller Faris margins and absence of formal significance tests warrant a measured interpretation.

**Figure 3.**
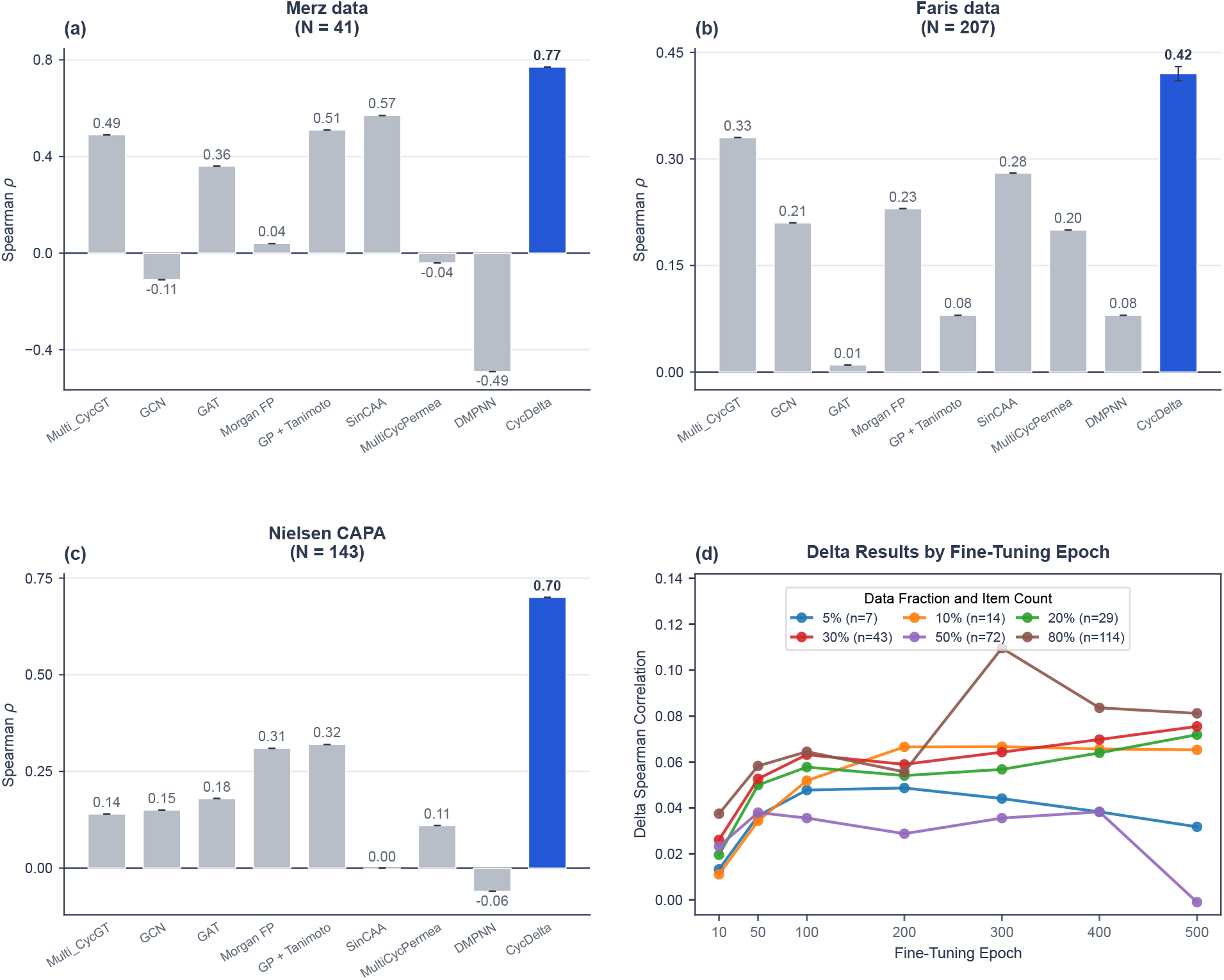
External transfer, unseen-assay transfer and few-shot adaptation. Spearman correlation is shown for (a) the Merz and (b) the Faris external data, using independent vertical scales. Error bars on CycDelta denote standard deviation over ten reference selections. (c) Spearman correlation for the prespecified zero-shot Nielsen CAPA evaluation (*N* = 143). Baseline methods are grey and CycDelta is blue. (d) Mean improvement in Nielsen test-set Spearman correlation relative to the frozen baseline, shown by labelled-data fraction and optimization epoch. The complete mean ± standard-deviation grid is reported in Supplementary Table S2. Higher values are better.

The contrast between the two external sets is informative. On the Merz data, GCN and MultiCycPermea were near zero and DMPNN was negatively correlated, indicating that success on the internal benchmark did not automatically transfer to this compact literature series. CycDelta’s high correlations and very small reference-selection variation suggest that its result was not driven by one favourable anchor. Performance was lower for every method on the larger Faris data, consistent with a more heterogeneous or difficult transfer setting. CycDelta nevertheless remained best on both correlation measures, so the external advantage was reproducible in direction.

### 4.4 Unseen-assay transfer on the Nielsen data

To examine transfer to an assay category absent from model development, we used the Nielsen data as a prespecified zero-shot evaluation. The assay vector was set to zero, and no Nielsen labels were used during fitting. CycDelta achieved Spearman *ρ* = 0.70, compared with 0.32 for GP + Tanimoto, the strongest baseline (Fig. 3c). This 0.38 margin supports transfer to an assay category absent from model development.

The Nielsen results reinforce the pattern observed on the external literature sets: strong internal absolute-value performance did not ensure transfer to a new data set. DMPNN, which led OD absolute-value Spearman correlation and MAE, fell to a Spearman correlation of −0.06 on Nielsen, while MultiCycPermea reached only 0.11. SinCAA obtained 0.00 on Nielsen despite reaching 0.57 on the Merz data, indicating that its external ranking performance was data-set dependent. Together with DMPNN’s negative correlations on the Merz data, these losses suggest that absolute-value models may rely partly on source-, scaffold- or measurement-specific property distributions that do not preserve peptide rankings after transfer. Morgan FP and GP + Tanimoto were more robust on Nielsen, but their correlations of 0.31 and 0.32 remained well below CycDelta’s 0.70, supporting direct within-source training as a route to more transferable comparative representations.

The assay-token analysis (Supplementary Table S1) indicates that conditioning on a mismatched permeability assay can reduce transfer correlation. For a new assay not represented during training, we therefore recommend using the unknown assay input (the zero vector) rather than substituting a token from a previously seen assay.

### 4.5 Few-shot fine-tuning on the Nielsen data

To determine whether wet-lab measurements from a new cyclic-peptide series could be fed back into CycDelta to improve predictions for other peptides in that series, we evaluated head-only fine-tuning across several labelled-data fractions. A subset of the Nielsen CAPA measurements was used to fine-tune the output head, and performance was evaluated on the remaining peptides from the same series. Fine-tuning generally produced positive improvements in test-set Spearman correlation, but neither increasing the labelled fraction nor extending optimization gave monotonic gains (Fig. 3d; Supplementary Table S2). With 5% of the data, improvement peaked at 0.049 at epoch 200 and then declined. The 10% condition reached a stable plateau from epochs 200–500, with a maximum of 0.067 at epoch 300. These results indicate that a small number of series-specific experimental measurements can fine-tune the output head and improve predictions for held-out peptides from the same series without changing the molecular encoder.

The 20% and 30% conditions continued improving through epoch 500, reaching 0.072 and 0.076, respectively. The 20% setting used only 29 labelled peptides and combined a competitive mean with lower variation than the 30% setting, making it a useful label-efficient operating point. However, because epochs were compared on the reported test splits, this observation should guide future validation rather than be treated as an independently selected optimum.

Larger fitting sets were not uniformly better. The 50% condition peaked at only 0.038 at epoch 400 and fell to −0.001 at epoch 500. The largest mean improvement, 0.11, occurred for 80% at epoch 300, but its standard deviation was 69% of the mean and the curve declined thereafter. This instability may reflect the small residual test set, split composition, reference choice or overfitting of the output head. Fraction-specific validation and uncertainty reporting are therefore more important than selecting one global epoch.

In summary, the present results suggest that measuring approximately 15–30 representative peptides and fine-tuning only the output head for roughly 200–300 epochs may provide an initial setting for a new cyclic-peptide series. These values should be regarded as provisional guidance rather than fixed requirements and adjusted using independent validation whenever possible.

### 4.6 Ablation

To identify which elements of CycDelta drive its performance, we ablated the training objective and individual representation components (Table 4). Removing the DMPNN caused the largest deterioration on OD within-source differences: Pearson/Spearman fell from 0.69/0.67 to 0.46/0.44 and MAE increased from 0.74 to 0.97. Removing within-source pair training reduced correlations to 0.59/0.57 and raised MAE to 0.86. Uni-Mol, RDKit and assay inputs made smaller complementary contributions.

**Table 4.** CycDelta ablation on OD within-source Δ and absolute-value evaluation.

| Variant | Within-source $\Delta$ | | | Absolute value | | |
| --- | --- | --- | --- | --- | --- | --- |
|  | Pearson | Spearman | MAE | Pearson | Spearman | MAE |
| Full CycDelta | 0.69±0.11 | 0.67±0.11 | 0.74±0.14 | 0.53±0.05 | 0.48±0.05 | 0.88±0.20 |
| No within-source pair training | 0.59±0.15 | 0.57±0.13 | 0.86±0.20 | 0.63±0.00 | 0.62±0.00 | 0.66±0.00 |
| No DMPNN | 0.46±0.16 | 0.44±0.16 | 0.97±0.18 | 0.25±0.07 | 0.22±0.08 | 1.26±0.34 |
| No assay method | 0.63±0.12 | 0.60±0.13 | 0.79±0.12 | 0.48±0.03 | 0.46±0.03 | 0.93±0.47 |
| No RDKit descriptors | 0.64±0.11 | 0.63±0.09 | 0.79±0.16 | 0.47±0.04 | 0.42±0.04 | 1.01±0.45 |
| No Uni-Mol | 0.62±0.09 | 0.60±0.07 | 0.78±0.11 | 0.45±0.04 | 0.46±0.05 | 1.02±0.52 |

The objective created a clear trade-off. Without within-source pair training, absolute-value Pearson, Spearman and MAE improved from 0.53/0.48/0.88 to 0.63/0.62/0.66, while pairwise performance declined. This interaction reinforces the need for a benchmark endpoint aligned with the intended comparative use.

Among the auxiliary feature channels, removing Uni-Mol caused the largest Pearson loss on within-source differences (0.07) and tied removal of the assay token for the largest Spearman loss (0.07). Removing RDKit descriptors reduced Pearson/Spearman by 0.05/0.04, while removing assay identity reduced them by 0.06/0.07. The corresponding MAE increases were modest compared with removal of the DMPNN. These patterns indicate complementary roles: message passing provides the dominant relational inductive bias, pretrained monomer embeddings encode structural context, explicit descriptors expose physicochemical quantities, and assay identity adjusts the mapping across measurement systems.

The absolute-value columns qualify the interpretation of the pairwise objective. The no-pair-training variant was best among the ablations on all three absolute metrics, whereas the full model was best on all three within-source metrics. Pairwise training should therefore be understood as task specialization rather than a universally beneficial regularizer. The ablation supports both the benchmark’s distinction between endpoints and the decision to evaluate CycDelta under both views.

## 5 From residue-level attribution to peptide design

CycDelta uses its differentiable residue representation for two connected purposes: explaining which residues support a pairwise permeability prediction and identifying substitutions that may shift permeability in a desired direction. These aims reflect the broader use of explainable machine learning to connect molecular predictions with testable chemical hypotheses [44]. The first is addressed through residue-level Integrated Gradients, while the second converts local feature sensitivity into candidate designs followed by full-model reranking.

### 5.1 Integrated-gradient attribution

To relate a predicted permeability change to individual residues, we applied Integrated Gradients (IG) to the residue states produced by the D-MPNN immediately before sum pooling [45]. Let 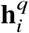 and 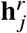 denote the post-D-MPNN states of query residue *i* and reference residue *j*, respectively. The part of the paired predictor downstream of these states is

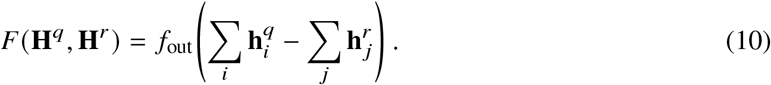

Using zero states for both peptides as the baseline, the signed attribution for residue *i* in peptide *m* ∈ {*q, r*} is

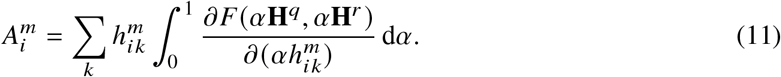

The integral was evaluated with 128 equal intervals using the trapezoidal rule. Across the evaluated test pairs, the maximum absolute completeness residual,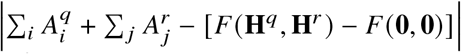 0.007. Positive 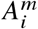 increases the predicted query-minus-reference permeability change and negative 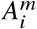 decreases it. Because the reference states enter Eq. (10) with a minus sign, the maps report molecule-centric values: 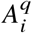 for a query residue and 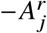 for a reference residue. Under this single convention a negative annotation denotes hindrance and a positive annotation denotes promotion of permeation in both columns, and colors carry the same sign. Values are shown without attribution normalization, and colors are scaled within each pair using the largest absolute residue attribution.

Reading a row is therefore additive. The predicted change is recovered as the sum of the query annotations minus the sum of the reference annotations. When the two peptides share a residue numbering, the difference between the query and reference annotations at one position gives that position’s contribution to the predicted change, and these position-wise differences sum to the same total. For peptides that differ throughout, the numbering is not shared and only the column-wise patterns are comparable.

Three examples illustrate the structural patterns exposed by the residue maps. In the first pair, CycDelta predicted a permeability change of −1.46, close to the measured change of −1.65 (Fig. 4a). The query residues all received negative contributions, with the largest magnitude at R1 (−0.63), followed by R3 (−0.49) and R2 (−0.44), while the reference annotations stayed close to zero (−0.17 to +0.04), so the predicted decrease is carried almost entirely by the query. The query contains polar and heteroatom-rich functionality, including a terminal urea-containing side chain, and the attribution pattern is consistent with the model treating these features as unfavourable in this molecular context. Polarity, hydrogen-bond exposure, lipophilicity and conformational shielding are established, interdependent determinants of cyclic-peptide permeability [24, 46–49]. This example shows that the model can localize a strongly negative prediction to chemically interpretable polar regions, although it does not establish polarity as an isolated causal determinant.

**Figure 4.**
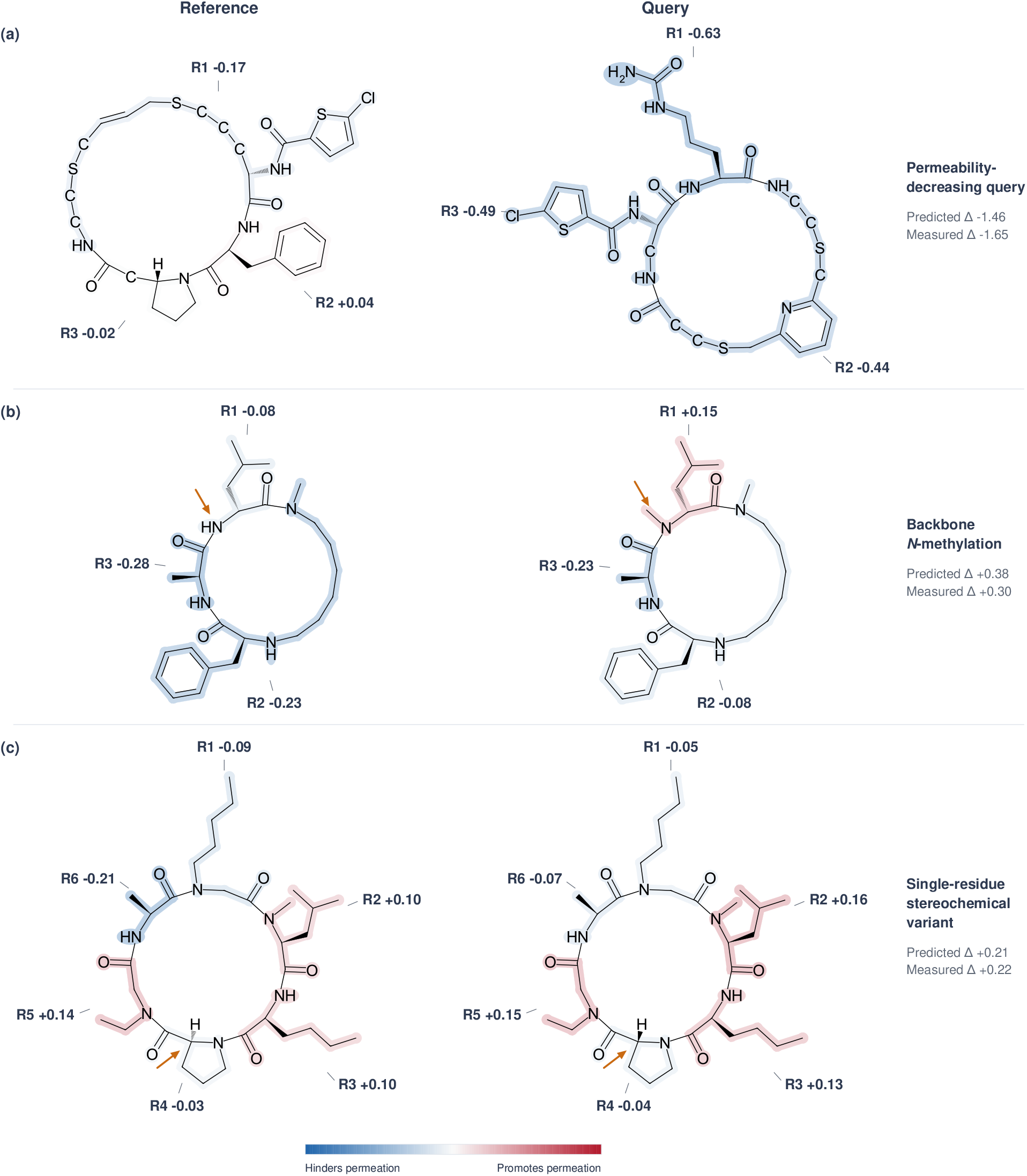
Residue-level IG maps for three representative permeability changes. Reference and query structures are shown in the left and right columns, respectively. (a) A permeability-decreasing query (predicted Δ = −1.46; measured Δ = −1.65). (b) Backbone *N*-methylation of R1 (predicted Δ = 0.38; measured Δ = 0.30). (c) A single-residue stereochemical variant at R4 (predicted Δ = 0.21; measured Δ = 0.22). In (b) and (c), where the two peptides differ at a single residue, the two structures share one depiction and one residue numbering wherever their chemistry agrees, so they can be compared position by position, and amber arrows mark the edited site in both columns; the peptides in (a) differ throughout and carry no arrow. Residue annotations are molecule-centric IG contributions, 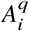 for query residues and 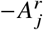 for reference residues, so that in both columns blue and negative denote hindrance of permeation while red and positive denote promotion, and the predicted change is the sum of the query annotations minus the sum of the reference annotations. Colors are scaled separately within each pair.

The second pair isolates an *N*-methylation that increased permeability (Fig. 4b). The query differs from the reference by a single backbone *N*-methyl group at the R1 D-leucine, a modification that removes one amide hydrogen-bond donor. Regioselective *N*-methylation can enhance permeability by altering hydrogen-bond exposure and conformational preferences, although its effect depends strongly on position and scaffold [6, 50, 51]. CycDelta predicted a change of 0.38, matching the measured increase of 0.30 in direction. The position-wise differences are +0.22 at the methylated R1, +0.14 at R2 and +0.05 at R3. Slightly more than half of the predicted gain is therefore assigned to the edited residue and the remainder is spread over the rest of the ring, indicating that the model represents *N*-methylation partly through its effect on the whole macrocycle rather than assigning the full gain to the methylated residue alone. This example shows that CycDelta can recover a chemically established permeability-enhancing modification while still treating it in a residue-graph context.

The third example isolates a stereochemical change at a proline residue. Stereochemical changes can reorganize intramolecular hydrogen bonds and alter permeability even when composition is unchanged [5, 51]. Conversion of the R4 proline configuration produced a measured permeability increase of 0.22, and CycDelta predicted a closely matching increase of 0.21 (Fig. 4c). Although the chemical edit is localized at R4, that position contributes almost nothing to the predicted change (−0.01), and the largest position-wise differences appear at other residues, particularly R6 (+0.14) and R2 (+0.06). This redistribution is consistent with the residue-graph encoder representing a stereochemical edit through its effect on the context of the whole macrocycle rather than assigning the full change to the edited residue alone.

Together, these examples show that CycDelta can recover the direction and approximate magnitude of selected permeability changes while providing residue-level accounts of its predictions. The maps should nevertheless be interpreted as explanations of model behaviour, not as proof of a general physicochemical mechanism; visually plausible attributions require model-sensitive checks and chemical validation [44, 52]. Establishing causal effects of polarity, *N*-methylation or stereochemistry will require controlled matched-series analyses and prospective experimental validation.

### 5.2 Gradient-guided monomer substitution

The differentiable encoder also supports candidate generation from one cyclic peptide, connecting differentiable molecular representations with property-directed search [23, 53]. The starting peptide is placed in both branches, giving an identity-pair prediction near 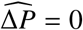, and gradients are calculated with respect to the query residue features. Replacing residue *i* by candidate monomer *c* receives the first-order score

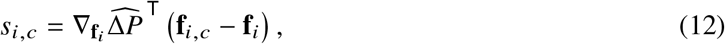

where **f**_*i*_ concatenates residue identity/modification, glycine-relative Uni-Mol and normalized RDKit features. The sign is reversed when decreased permeability is requested.

This score is used for screening rather than as the final prediction. Candidate monomers may be restricted by class or training frequency; proposed substitutions are assembled into cyclic structures and rejected if chemical validation fails, reflecting the need to constrain generated structures to chemically interpretable space [23, 28]. The highest-scoring valid candidates are then evaluated by a complete CycDelta forward pass and ranked by the recomputed permeability change. The procedure therefore does not differentiate through a discrete SMILES string; it uses local feature gradients to prioritize substitutions and verifies them with the full model.

We applied this procedure to one cyclic-peptide reference while restricting the candidate library to natural L-amino acids. The ten highest-ranked candidate entries all produced positive full-model predictions, ranging from 0.79 to 1.31, and concentrated at two positions. Seven entries targeted the R2 histidine and represented six amino-acid substitutions: H-to-Ile, H-to-Leu, H-to-Pro, H-to-Phe, H-to-Val and H-to-Gly. The highest predicted changes were obtained for H-to-Ile (1.31) and H-to-Leu (1.19). Three entries targeted the R1 threonine and represented T-to-Ile and T-to-Leu, with predicted changes of up to 0.88 and 0.87, respectively.

This ranking reveals a consistent design pattern. At R2, most high-ranking substitutions replace the potentially ionizable, hydrogen-bonding imidazole of histidine with less polar aliphatic, aromatic or conformationally restricted side chains. At R1, replacing threonine with Ile or Leu removes the hydroxyl group and increases hydrophobic character. These trends are compatible with established permeability determinants, but increasing lipophilicity can also reduce aqueous solubility and must be interpreted in a scaffold-dependent manner [24, 48, 49]. The appearance of Gly among the R2 candidates further indicates that the model considers side-chain size and conformational effects in addition to hydrophobicity. R2 accounted for seven of the top ten suggestions and was therefore the dominant optimization hotspot, while R1 provided a second independently modifiable site. The highest-ranked R2 substitution and the highest-ranked R1 substitution are shown in Fig. 5a and 5b, respectively.

**Figure 5.**
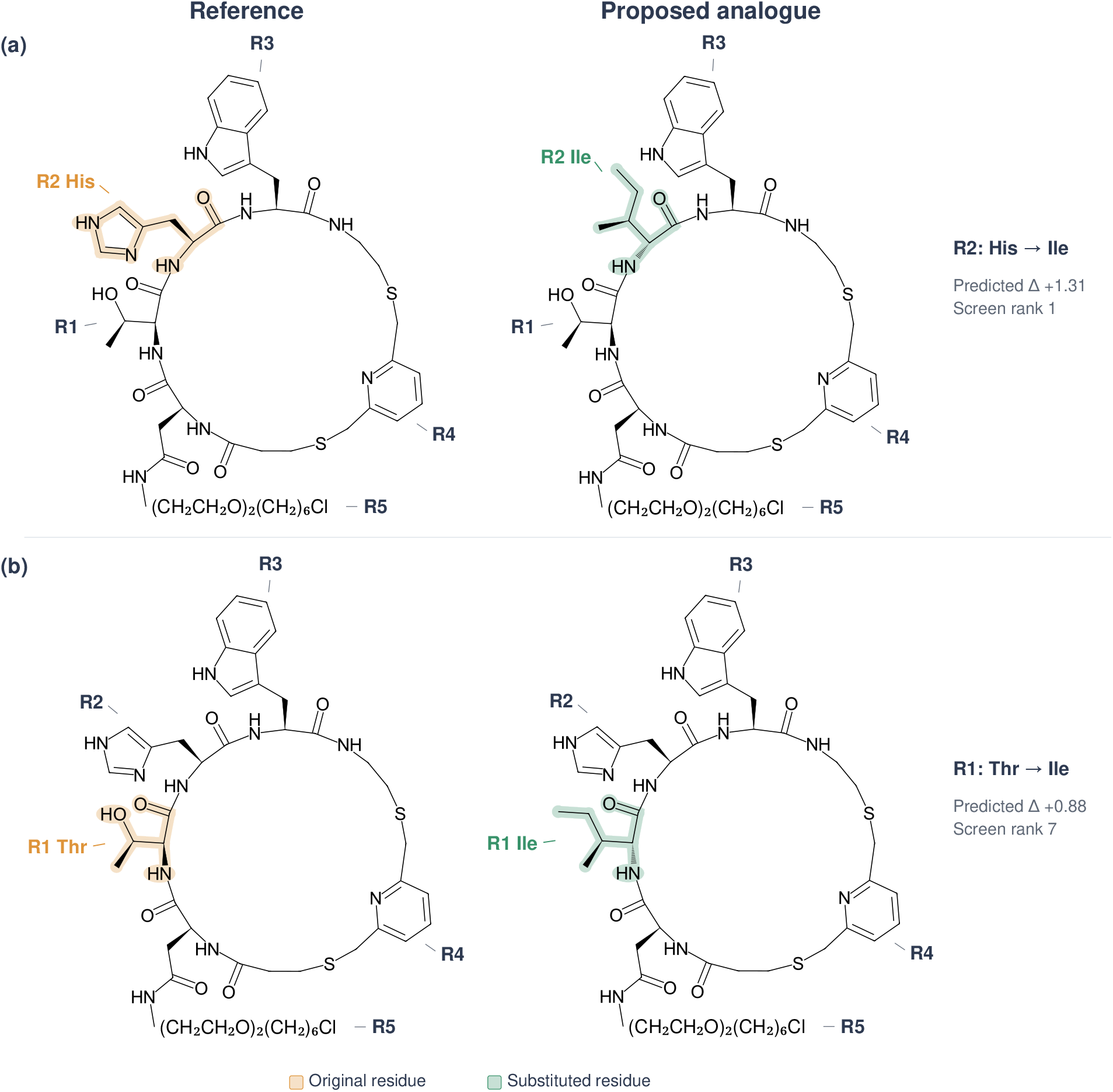
Gradient-guided monomer substitutions proposed for one measured lead. Both proposals were generated from the same cyclic-peptide reference (left column) and are shown next to the corresponding analogue (right column); the substituted residue is tinted and every other residue keeps the reference geometry, so the two structures in a row differ only where the chemistry differs. (a) The highest-ranked proposal replaced histidine at R2 with isoleucine (predicted Δ = 1.31). (b) The seventh-ranked proposal replaced threonine at R1 with isoleucine (predicted Δ = 0.88). The poly(ethylene glycol)–chloroalkyl tail carried by the R5 side-chain amide is drawn in condensed form. Predicted values are query-minus-reference permeability changes from full-model reevaluation, and the two proposals identify independently modifiable positions in the same reference.

### 5.3 Application scope

CycDelta is intended for applications in which a team has measured a cyclic-peptide lead and must prioritize candidates for the next experimental round. The model uses the measured lead as an anchor to rank proposed analogues and estimate the direction of change without requiring perfect calibration on the absolute assay scale, paralleling the comparative logic of matched-pair medicinal chemistry and model-guided analogue generation [19, 23]. This use is especially attractive when source-associated offsets are comparable to the molecular effect being optimized.

Although CycDelta supports local analogue prioritization, its comparative prediction task is broader than single-residue modification. Sources may contain close analogue series, but they may also span wider structural diversity; no similarity threshold or single-residue constraint is imposed. The gradient-guided workflow focuses the general model on design by proposing chemically valid, position-specific substitutions from a measured lead. The ranked His-to-Ile and Thr-to-Ile examples demonstrate this operational use and provide concrete candidates for synthesis and experimental testing.

## 6 Conclusion

CycDelta demonstrates the value of aligning model architecture with the comparative endpoint. Its shared multimodal residue-graph encoder and direct training on within-source differences achieved Pearson/Spearman correlations of 0.89/0.87 in the ID setting and 0.69/0.67 in the primary scaffold-OD setting. On OD data, the correlation margin over the strongest baseline was 0.23 for both metrics, showing that the model preserved permeability ordering and covariation across unseen scaffold groups. The results also establish that pairwise prediction is a distinct modelling objective: performance on absolute permeability alone did not reliably predict performance on within-source changes.

CycDeltaBench provides a controlled evaluation framework for this comparative formulation by changing the unit of permeability modelling from an isolated cyclic peptide to a comparison made within a common literature or experimental source. This setup aligns evaluation with a central design question: given a measured reference peptide, which candidate is likely to improve permeability, and by how much? Fixed data curation, pair construction, reference selection, ID and scaffold-OD partitions, external tests and metrics enable consistent comparison of CycDelta with alternative methods.

The transfer results extend this advantage beyond the internal benchmark. CycDelta achieved Pearson/Spearman correlations of 0.74/0.77 on the Merz data and 0.42/0.42 on the Faris data, leading all evaluated methods on both external sets. In zero-shot transfer to the unseen Nielsen CAPA assay, it reached a Spearman correlation of 0.70 compared with 0.32 for the strongest baseline. Head-only fine-tuning further showed that a limited number of measurements from a new peptide series can improve predictions while preserving the pretrained molecular encoder.

Residue-level IG maps connect these predictive results to chemically interpretable patterns involving polarity, backbone *N*-methylation and stereochemistry. The same differentiable representation supports gradient-guided monomer screening, chemical-validity filtering and full-model reranking, providing a practical route from a measured lead to prioritized analogue hypotheses. These capabilities position CycDelta as both a comparative prediction model and a design-support framework for cyclic-peptide optimization. Some published predictors, however, require specialized sequence, conformational or image inputs rather than a SMILES string, which can limit their convenience in routine application.

The current study uses literature source as the available experimental context and will benefit from future prospective, assay-matched analogue campaigns. Nevertheless, the consistent gains across scaffold-OD, external and unseen-assay evaluations support the central conclusion: learning permeability changes directly within source produces transferable comparative representations and provides a useful bridge between benchmark evaluation and cyclic-peptide design.

## Supplementary information

Supplementary Fig. S1 shows the agreement of PAMPA measurements for identical structures reported across literature sources. Supplementary Figs. S2 and S3 show observed-versus-predicted scatter and regression plots for CycDelta at reference-selection seed 0 in the scaffold-OD and ID settings. Supplementary Table S1 reports exploratory assay-token sensitivity on the Nielsen data, and Supplementary Table S2 reports the head-only fine-tuning grid.

## Author contributions

J.Q. implemented the model, carried out the training and evaluation experiments and wrote the manuscript. Y.Z. contributed to the collection of part of the evaluation data. Y.C. and Z.G. contributed to the study design and provided advice on the modelling approach and the evaluation strategy. L.W. and S.L. contributed to the cyclic-peptide chemistry perspective, the interpretation of the results and the discussion of design applications. D.W. and M.Y. conceived the study, supervised the work and provided critical revisions of the manuscript. All authors read and approved the final manuscript.

## Data availability

The development records were curated from CycPeptMPDB v1.2 [7], which is publicly available, and the external evaluation sets were derived from the previously reported Faris [34], Merz [35] and Nielsen [36] studies. The curated CycDeltaBench records, partitions and query–reference pair definitions are distributed together with the source code at https://github.com/cyclontx/CycDelta.

## Code availability

The source code for CycDelta, together with the CycDeltaBench data-curation, pair-construction and evaluation pipeline, is openly available at https://github.com/cyclontx/CycDelta.

## Ethics approval

Not applicable; this study used previously reported molecular and assay data.

## Supplementary Figures and Tables

**Supplementary Fig. S1.**
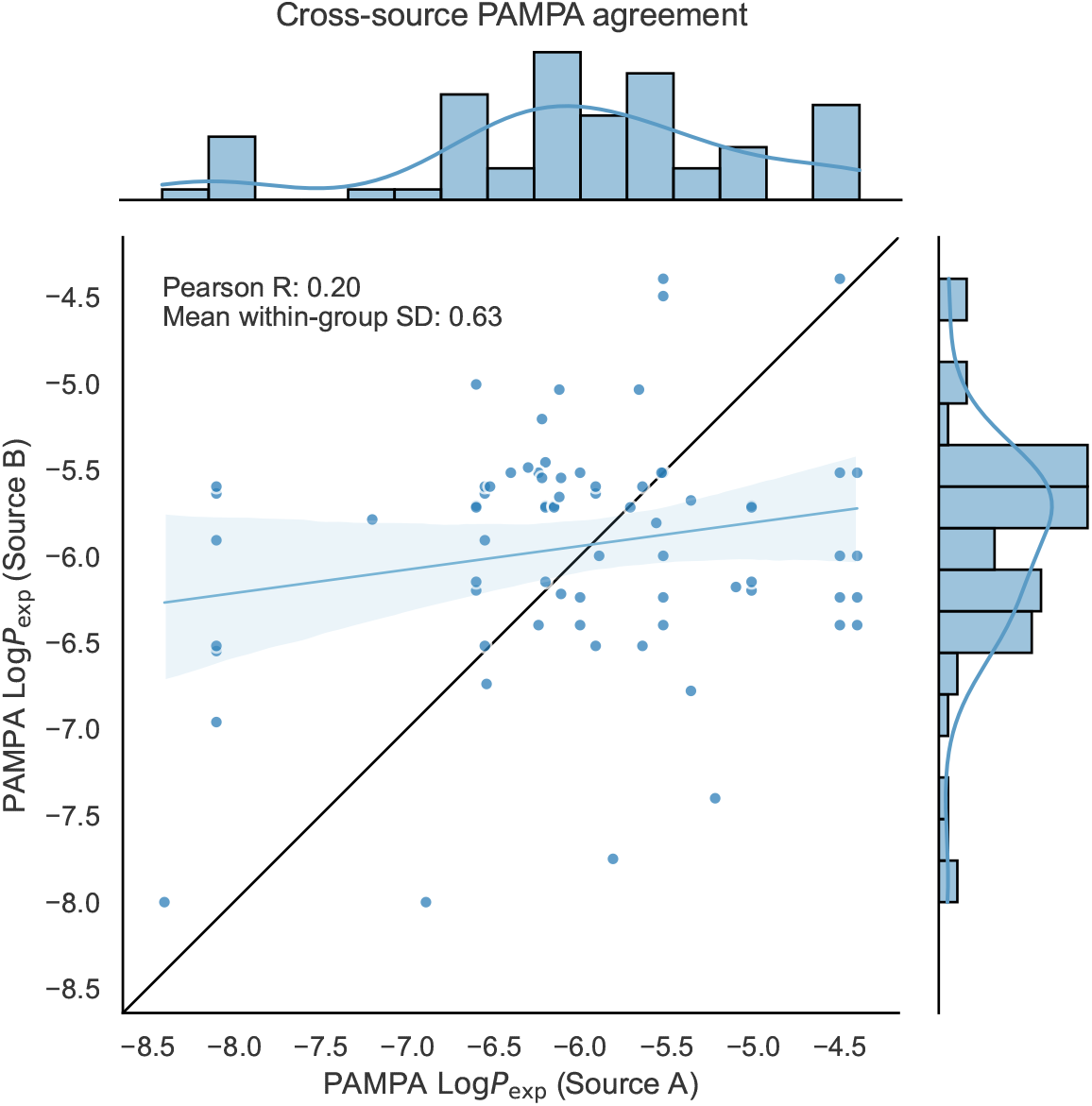
Agreement of PAMPA permeability measurements across literature sources. Each point represents one pairwise comparison between measurements of the same peptide structure reported in two different sources. The 21 matched structures comprised 57 measurements and generated 73 unique source pairs; structures reported by more than two sources therefore contribute multiple, non-independent points. The black diagonal denotes identity, the blue line and shaded band show the fitted linear regression and its 95% confidence interval, and the marginal distributions are shown above and to the right. Pearson correlation and the unweighted mean of the 21 within-structure sample standard deviations are displayed in the plot.

**Supplementary Fig. S2.**
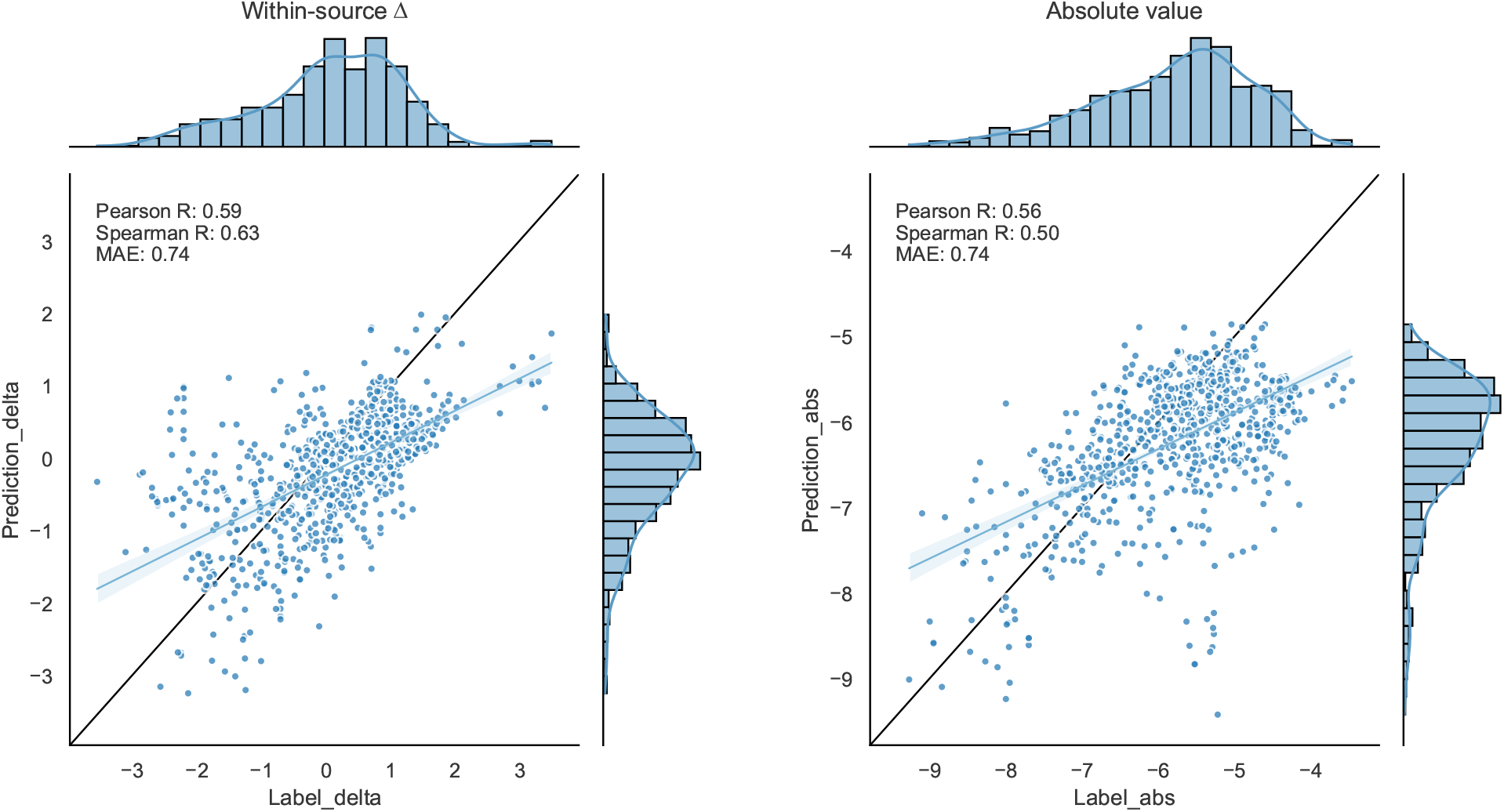
CycDelta predictions in the scaffold-OD setting for reference-selection seed 0. Observed values are plotted against model predictions for within-source permeability differences (left) and reconstructed absolute permeability values (right). The blue line and shaded band show the fitted linear regression and its confidence interval; the black diagonal denotes identity. Marginal distributions are shown above and to the right of each scatter plot.

**Supplementary Fig. S3.**
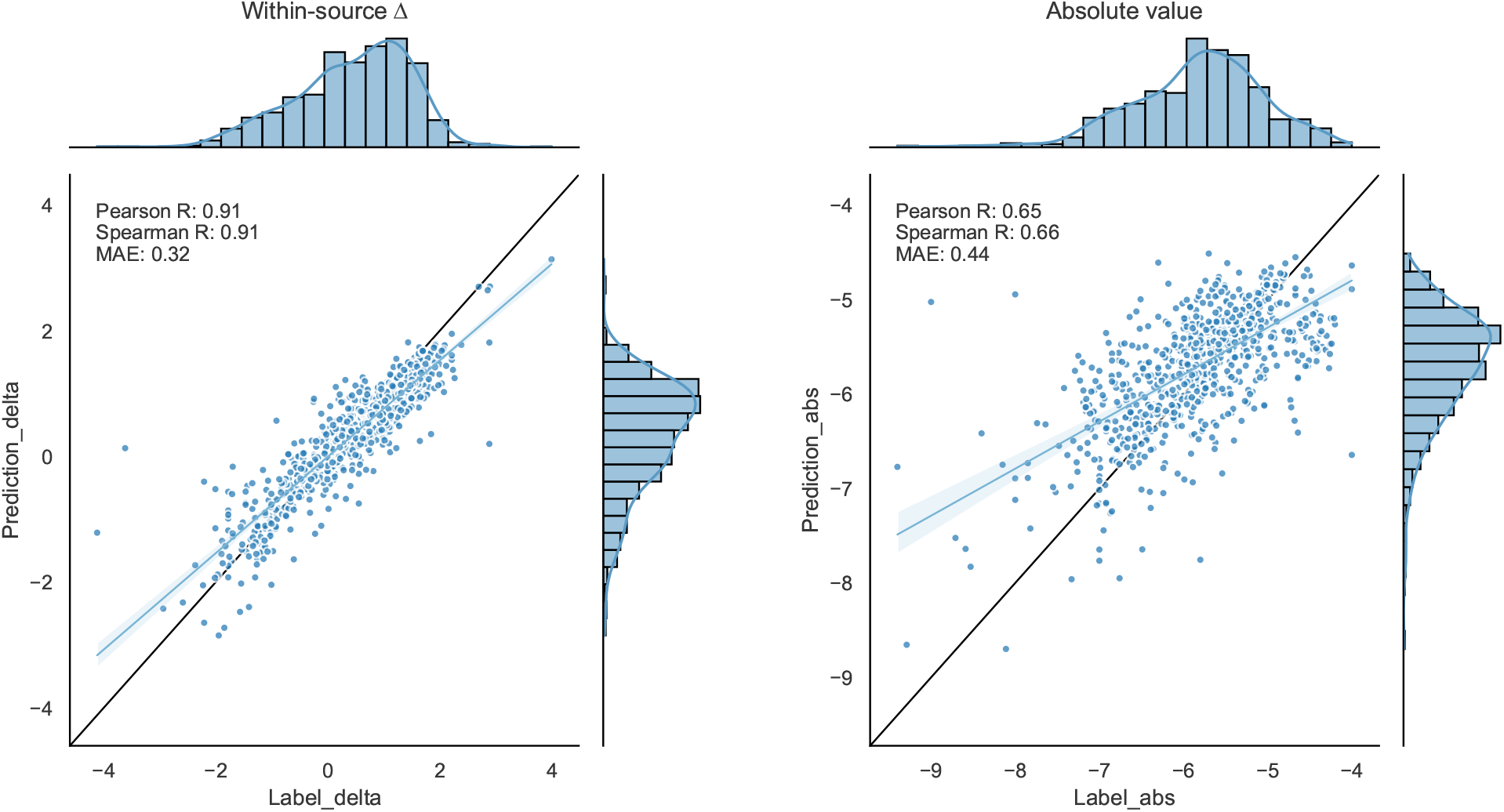
CycDelta predictions in the ID setting for reference-selection seed 0. Observed values are plotted against model predictions for within-source permeability differences (left) and reconstructed absolute permeability values (right). The blue line and shaded band show the fitted linear regression and its confidence interval; the black diagonal denotes identity. Marginal distributions are shown above and to the right of each scatter plot.

**Supplementary Table S1.** Exploratory assay-token sensitivity of CycDelta on the Nielsen data (*N* = 143). Bold identifies the prespecified unknown-token result, not the post hoc numerical maximum.

| Assay token | Spearman $\rho$ |
| --- | --- |
| Unknown (zero vector) | **0.70** |
| PAMPA | 0.69 |
| CACO2 | 0.69 |
| MDCK | 0.61 |
| RRCK | 0.71 |

**Supplementary Table S2.** Complete head-only fine-tuning results on the Nielsen data. Each entry is the mean improvement in test-set Spearman correlation over the frozen baseline on the same split; “±” denotes standard deviation across ten seeded splits and associated reference selections.

| Fraction | *N* | Epoch 10 | Epoch 50 | Epoch 100 | Epoch 200 | Epoch 300 | Epoch 400 | Epoch 500 |
| --- | --- | --- | --- | --- | --- | --- | --- | --- |
| 5% | 7 | 0.013±0.007 | 0.037±0.017 | 0.048±0.019 | 0.049±0.031 | 0.044±0.034 | 0.038±0.034 | 0.032±0.037 |
| 10% | 14 | 0.011±0.005 | 0.034±0.013 | 0.052±0.011 | 0.067±0.009 | 0.067±0.014 | 0.066±0.018 | 0.065±0.022 |
| 20% | 29 | 0.020±0.007 | 0.050±0.012 | 0.058±0.020 | 0.054±0.025 | 0.057±0.026 | 0.064±0.024 | 0.072±0.019 |
| 30% | 43 | 0.026±0.005 | 0.053±0.020 | 0.063±0.037 | 0.059±0.050 | 0.064±0.051 | 0.070±0.044 | 0.076±0.037 |
| 50% | 72 | 0.023±0.015 | 0.038±0.036 | 0.036±0.045 | 0.029±0.056 | 0.036±0.069 | 0.038±0.071 | -0.001±0.093 |
| 80% | 114 | 0.038±0.022 | 0.058±0.056 | 0.065±0.060 | 0.056±0.084 | 0.110±0.076 | 0.084±0.105 | 0.081±0.079 |

